# COMPASS: a COMprehensive Platform for smAll RNA-Seq data analySis

**DOI:** 10.1101/675777

**Authors:** Jiang Li, Alvin T. Kho, Robert P. Chase, Lorena Pantano-Rubino, Leanna Farnam, Sami S. Amr, Kelan G. Tantisira

## Abstract

**Background:** Circulating RNAs are potential disease biomarkers and their function is being actively investigated. Next generation sequencing (NGS) is a common means to interrogate the small RNA’ome or the full spectrum of small RNAs (<200 nucleotide length) of a biological system. A pivotal problem in NGS based small RNA analysis is identifying and quantifying the small RNA’ome constituent components. Most existing NGS data analysis tools focus on the microRNA component and a few other small RNA types like piRNA, snRNA and snoRNA. A comprehensive platform is needed to interrogate the full small RNA’ome, a prerequisite for down-stream data analysis.

**Results:** We present COMPASS, a comprehensive modular stand-alone platform for identifying and quantifying small RNAs from small RNA sequencing data. COMPASS contains prebuilt customizable standard RNA databases and sequence processing tools to enable turnkey basic small RNA analysis. We evaluated COMPASS against comparable existing tools on small RNA sequencing data set from serum samples of 12 healthy human controls, and COMPASS identified a greater diversity and abundance of small RNA molecules.

**Conclusion:** COMPASS is modular, stand-alone and integrates multiple customizable RNA databases and sequence processing tool and is distributed under the GNU General Public License free to non-commercial registered users at https://regepi.bwh.harvard.edu/circurna/ and the source code is available at https://github.com/cougarlj/COMPASS.

## Background

The human circulation system (serum and plasma) contains various types of RNA molecules, including fragmental mRNA, miRNA, piRNA, snRNA, snoRNA, and some other non-coding sequences [1, 2]. Studies have shown the biomarker potential of circulating RNAs in cancer [3], cardiovascular disease [4], and asthma [5]. Moreover, other types of DNA and RNA fragments discovered in the human circulating system have been implicated as potential causes of chronic disease [6, 7].

Small RNA sequencing (RNA-seq) technology was developed successfully based on special isolation methods and the RNA-seq technique, which facilitates the investigation of a comprehensive profile of circulating RNAs [8, 9]. In anticipation of a continued growing number of circulating RNAs studies, a comprehensive and stable platform is needed to identify the RNA classification, RNA read counts, differential expression between case and control samples, including both human and non-human (e.g. microbiome) small RNAs (<200 nucleotide length). Previous efforts to characterize small RNAs have focused primarily on microRNAs (miRNAs). For instance, sRNAnalyzer is a comprehensive and customizable pipeline for the small RNA-seq data centred on microRNA (miRNA) profiling [10]. sRNAtoolbox is a web-based small RNA research toolkit [11] and SeqCluster has started to focus on non-miRNAs by comparing the sequence similarity [12]. Some efforts have begun to characterize the full spectrum of small RNAs of a biological system (the small RNA’ome), such as Oasis2 and exceRpt. Oasis2 is a web server for small RNA-seq data analysis [13]. ExceRpt, maintained by the Extracellular RNA Communication Consortium (ERCC), is an extensive and commonly used web-based pipeline for extra-cellular RNA profiling [14]. Currently, Oasis2 does not support microbial RNA identification and exceRpt provides few microbiome annotations. Both the tools need users to upload the original sequencing files, which is un-workable for big data.

To profile intracellular and extracellular small RNA’omes through the small RNA-seq data, we built a comprehensive platform COMPASS to identify and quantify diverse RNA molecule types, including miRNA, piRNA, snRNA, snoRNA, tRNA, circRNA and the fragmental microbial RNA. COMPASS is built using Java and works as stand-alone providing detailed annotation for each type of small RNAs including microbial constituents. It currently uses STAR [15] and BLAST [16] for alignment and sequence comparison. It takes FASTQ file as inputs and outputs the counts profile of each type of RNA molecule type per FASTQ file (typically representing a sample). When case and control files are marked, COMPASS can perform a differential expression analysis with the p value from the Mann-Whitney U test as default.

## Implementation

COMPASS is built using Java and composed of five functionally independent and customizable modules: Quality Control (QC), Alignment, Annotation, Microbe and Function (see Fig 1). Users can run all the modules as an integrated pipeline or just use certain modules. Since COMPASS is a stand-alone platform, it can be installed in any desktop or server, which maximizes data security and bypasses time/effort transferring data offsite that web-based tools need.

**Fig. 1.**
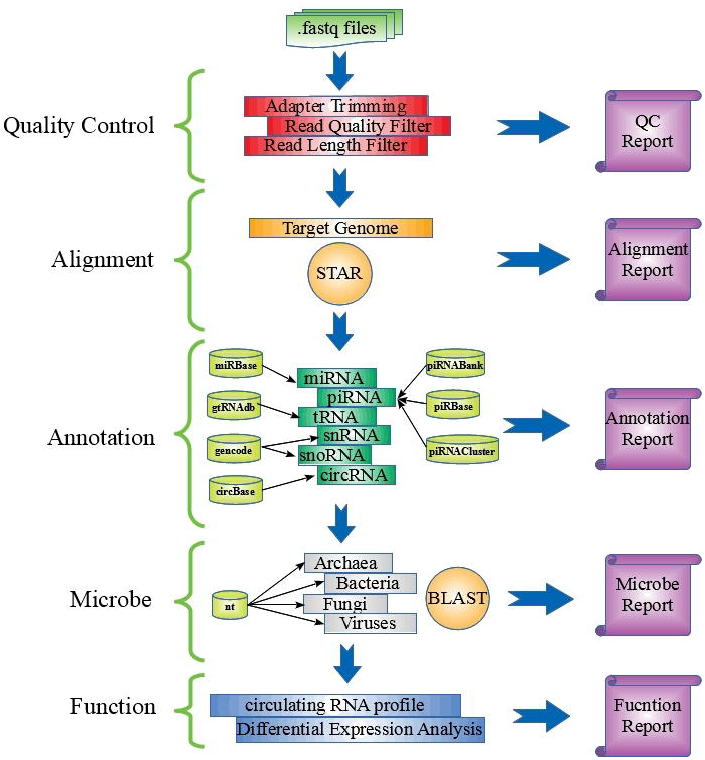
The structure of COMPASS platform. COMPASS is a comprehensive platform for circulating RNA analysis.

### 2.1 Quality Control (QC) Module

FASTQ files from the small RNA-seq of biological samples are the default input. First, the adapter portions of a read are trimmed along with any randomized bases at ligation junctions that are produced by some small RNA-seq kits (e.g., NEXTflexTM Small RNA-Seq kit) [17]. The read quality of the remaining sequence is evaluated using its corresponding PHRED score. Poor quality reads are removed according to quality control parameters set in the command line (-rh 20 –rt 20 –rr 20). Users can specify qualified reads of specific length intervals for input into subsequent modules.

### 2.2 Alignment Module

COMPASS uses STAR v2.5.3a [15] as its default RNA sequence aligner with default parameters which are customizable on the command line. Qualified reads from the QC module output are first mapped to the human genome hg19/hg38, and then aligned reads are quantified and annotated in the Annotation Module. Reads that could not be mapped to the human genome are saved into a FASTA file for input into the Microbe Module.

There are two scenarios where multi-aligned reads may exist when aligned against a reference genome. First, one small RNA read could be aligned to multiple distinct genomic locations. For example, the miRNA hsa-miR-1302 can derive from 11 potential pre-miRNAs. In this scenario, COMPASS will only count once with the multi-aligned read. Second, two or more distinct small RNAs could have overlapping sequences. For example, miRNA has-let-7a-5p (**UGAGGUAGUAGGUUGUAUAGUU**) and piRNA has_piR_008113 (**UGAGGUAGUAGGUUGUAUAGUU**UUAGGGUC) have significant sequence overlaps. In this case, each small RNA will be assigned with one count.

### 2.3 Annotation Module

COMPASS currently uses several different (and expandable) small RNA reference databases for annotating human genome mapped reads: miRBase [18] for miRNA; piRNABank [19], piRBase [20] and piRNACluster [21] for piRNA; gtRNAdb [22] for tRNA; GENCODE release 27 [23] for snRNA and snoRNA; circBase [24] for circular RNA. To harmonize the different reference human genome versions in these databases, we use an automatic LiftOver created by the UCSC Genome Browser Group. All the databases used are pre-built to enable speedy annotation. For each RNA molecule, COMPASS provides both the read count and indicates the database items that support its annotation. Using the command line parameter –abam, COMPASS will output all the reads that are annotated to a specific type of RNAs.

### 2.4 Microbe Module (Optional)

The qualified reads that were not mapped to the human genome in the Alignment Module are aligned to the nucleotide (nt) database [25] from UCSC using BLAST. Because of the homology between species, one read may be aligned to many species and COMPASS will list all the potential taxa with read count according to the phylogenetic tree as default. The four major microbial taxons archaea, bacteria, fungi and viruses are supported. To optimize processing the BLAST results, a fast accessing and parsing text algorithm is used [26].

### 2.5 Function Module

The read count of each RNA molecule that is identified in the Annotation Module is outputted as a tab-delimited text file according to RNA type. With more than one sample FASTQ file inputs, the output are further aggregated into a data matrix of RNA molecules as rows and samples as columns showing the read counts of an RNA molecule across different samples. The default normalization method is Count-per-Million (CpM), which normalizes each sample library size into one million reads. The user can mark each sample FASTQ file column as either a case or a control in the command line, and perform a case versus control differential expression analysis for each RNA molecule using the Mann-Whitney rank sum test (Wilcoxon Rank Sum Test) as the default statistical test.

## Results and discussion

We processed small RNA-seq FASTQ files from the serum of 12 healthy human subjects in a performance study through COMPASS to evaluate its performance on diverse types of RNA molecules, and compare it to a previously published web-based pipeline exceRpt [14]. Serum samples were prepared using NEXTflex Small RNA Kit and sequenced through the Illumina platform.

We run COMPASS on server with 30GB RAM. COMPASS takes ~15 minutes per sample per processor for the first three modules. More processing time is required if the microbiome module is required. The output files of each type of RNAs contain four columns: DB (databases used for annotation), Name (name of the RNA), ID (general id of the RNA) and Count (counts of reads). A summary count file including all samples can be obtained from the function module (-fun_merge).

The read length distribution of 12 serum samples was described in Fig 2. The length of raw reads was 50nt and after trimming adapters and 4 random bases at both 5’ and 3’ ends, the read length varied from 0nt to 42nt. In general, without size selection at the library preparation stage, each read length distribution of one sample has 4 peaks. The miRNAs should be located around the main peak at 22nt according to their structure characteristic. The piRNAs were distributed around 30nt and the 32nt peak represents the Y4-RNA (Ro60-associated Y4) and some tRNA fragments [27]. The 42nt (or trimmed maximum read length in this study) might represent snRNAs, mRNA fragments and microbial RNAs. The snoRNA was overlapped with miRNA in a great measure. In addition, there were still large part of short RNA fragment around 12nt, which may come from some RNA degradation products or even some unknown RNA classes.

**Fig. 2.**
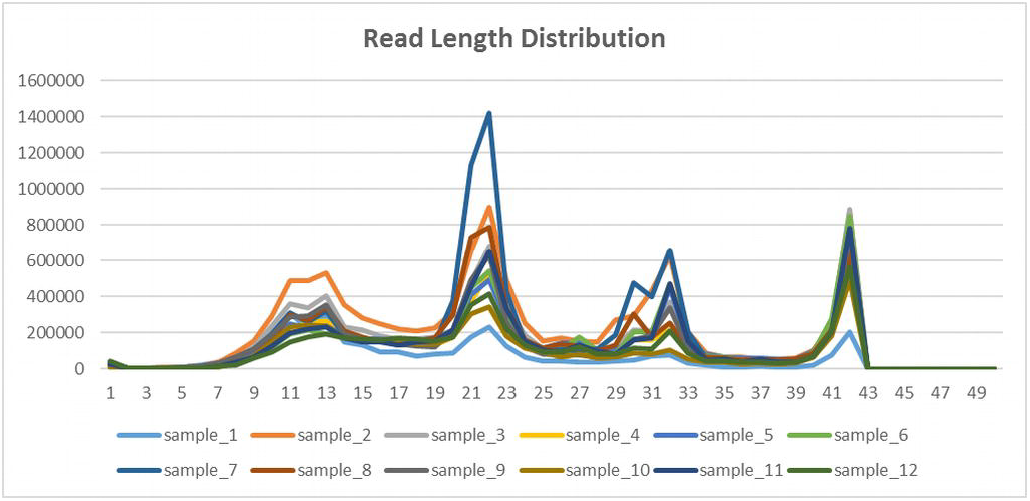
The read length distribution of 12 serum testing samples.

COMPASS identified diverse types of RNA molecules in this study including miRNAs, piRNAs, snRNAs, snoRNAs, tRNAs, circRNAs and RNAs in microbes (see Fig 3). We used a read count threshold of 5 to indicate that a RNA molecule was detected (≥5) or not (<5). In total, COMPASS detected 375 miRNAs, 280 piRNAs, 167 snRNAs, 88 snoRNAs, 401 tRNAs and 7285 circRNAs, as well as 608 archaea, 103825 bacteria, 45343 fungi and 208 viruses. The tRNAs were marked with the tRNAscan-SE IDs which were based on tRNA genes [28]. 7285 circRNAs were identified, which was much higher than other small RNAs. It might be because that the total number of circRNAs in human genome is huge. According to the statistics in CIRCpedia (v2), the human genome v38 (hg38) may contain 183,943 circRNAs [29]. The species of microbe were still large in number, which may be caused by the cross species homology. If one sequence read aligned to multiple homologous species, COMPASS will output all the species without bias.

**Fig. 3.**
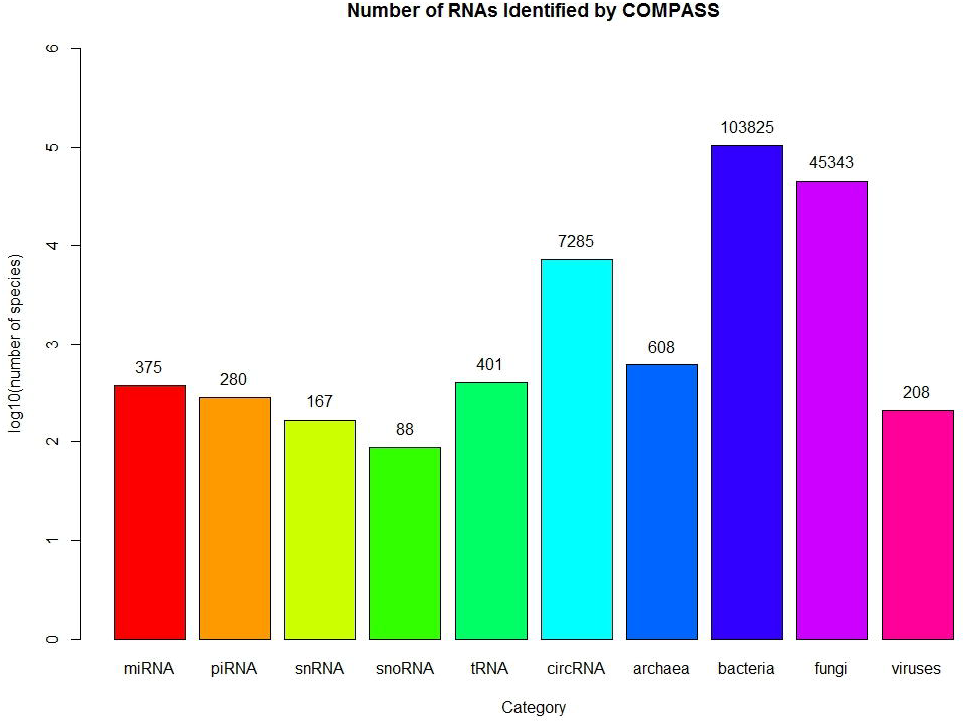
Number of RNAs Identified by COMPASS through 12 serum samples.

Compared to exceRpt outputs of the same study data (See Fig 4), COMPASS generally shared a large proportion of commonly identified small RNAs with COMPASS identifying more unique RNAs than exceRpt. For miRNAs, both COMPASS and exceRpt identified 358 (90% of total miRNAs) miRNAs. Although exceRpt had 24 unique miRNAs, 18 (75%) of them were only detected in one sample. We listed the comparison of all the 12 samples between COMPASS and exceRpt in Table 1. In each sample, the median counts of miRNAs from COMPASS and exceRpt are nearly the same. COMPASS and exceRpt had 27 common snoRNAs, among which 11 (41%) of them were detected only in one sample by COMPASS and 15 (56%) of them detected only in one sample by exceRpt. COMPASS had 61 unique snoRNAs, of which 41 (67%) snoRNAs existed only in one sample. However, exceRpt had 39 unique snoRNAs but 32 (82%) of them existed in one sample. snoRNAs were stable in circulation and they have been validated as biomarkers in some disease studies [30, 31]. Compared with exceRpt, COMPASS may have a more robust results in snoRNAs detection. In the comparison of tRNAs, we reclassified tRNAs according to the amino acid it carries as exceRpt did.

**Table 1.** miRNAs identified by COMPASS and exceRpt among each sample.

|  | miRNA |  |  |  |  |
| --- | --- | --- | --- | --- | --- |
|  | COMPASS<br>(median) | exceRpt<br>(median) | Overlap | COMPASS_Unique | exceRpt_Unique |
| SAMPLE_1 | 64(875.5) | 66(898) | 63 | 1 | 3 |
| SAMPLE_2 | 163(491) | 170(492.5) | 159 | 4 | 11 |
| SAMPLE_3 | 174(333) | 176(337) | 164 | 10 | 12 |
| SAMPLE_4 | 147(545) | 148(557.5) | 140 | 7 | 8 |
| SAMPLE_5 | 123(644) | 123(699) | 117 | 6 | 6 |
| SAMPLE_6 | 104(723.5) | 102(754.5) | 98 | 6 | 4 |
| SAMPLE_7 | 240(400.5) | 249(397) | 234 | 6 | 15 |
| SAMPLE_8 | 153(607) | 156(568.5) | 150 | 3 | 6 |
| SAMPLE_9 | 195(343) | 200(348) | 187 | 8 | 13 |
| SAMPLE_10 | 91(767) | 88(747) | 87 | 4 | 1 |
| SAMPLE_11 | 125(668) | 126(677.5) | 121 | 4 | 5 |
| SAMPLE_12 | 108(671) | 111(703) | 104 | 4 | 7 |

**Fig. 4.**
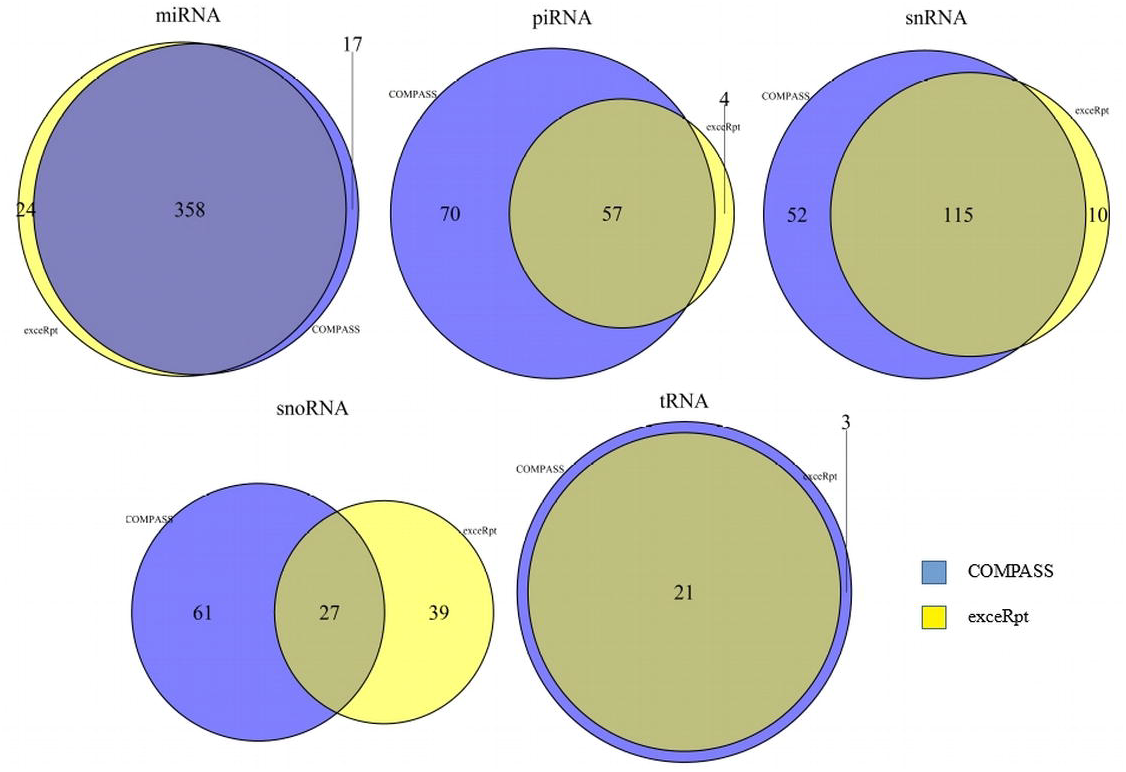
Summarize the comparison between COMPASS and exceRpt.

The comparisons of piRNAs, snRNAs, snoRNAs and tRNAs at the sample level were shown in **Table S1-S4**. COMPASS can always identify more piRNAs than exceRpt (**Table S1**). The reason may be that COMPASS use not only piRNABank database but also piRBase to annotate piRNAs. For snRNAs (**Table S2**), snoRNAs (**Table S3**) and tRNAs (**Table S4**), COMPASS and exceRpt can detect a large set of common RNAs. There were more COMPASS unique RNAs than than exceRpt unique RNAs, and a greater proportion of COMPASS unique RNAs were detected in 2 or more samples than exceRpt unique RNAs. The median read count values from COMPASS is usually larger than exceRpt. This could be because COMPASS outputs the total read count for each RNA, while exceRpt normalizes the count by copy numbers. This will significantly decrease the count number when the RNA has more copies in the genome. In addition, exceRpt annotates the RNA types in order of priority (miRNA>tRNA>piRNA>snRNA and snoRNA>circRNA), so that when an aligned read has been annotated to a certain small RNA type, the read will not be annotated to the other types at a lower priority order. COMPASS annotates an aligned read to all RNA types without an order of priority.

## Conclusions

COMPASS is a comprehensive modular stand-alone platform for the small RNA-seq data analysis. As a stand-alone platform, it bypasses data transfer effort/time/risk offsite that web-based tools need.

Its modularity allows the user to run all modules together as a complete basic small RNA analysis pipeline or specific modules as needed. Its pre-built RNA databases and sequence read processing tools enable turnkey basic small RNA analysis from identification, quantification to basic differential analysis. These pre-built databases/tools are customizable and expandable. COMPASS is distributed under the GNU General Public License free to non-commercial registered users at https://regepi.bwh.harvard.edu/circurna/ and the source code is available at https://github.com/cougarlj/COMPASS.

## Availability and requirements

Project name: COMPASS

Project home page: https://regepi.bwh.harvard.edu/circurna/

Operating System(s): Linux, Mac and Windows

Programming language: Java

Other requirements: Java 1.7 or higher

License: GNU GPL

Any restrictions to use by non-academics: licence needed

## Abbreviations

circRNA: circular RNA
COMPASS: a COMprehensive Platform for smAll RNA-Seq data analysis
exceRpt: The extra-cellular RNA processing toolkit
miRNA: microRNA
NGS: Next Generation Sequencing
piRNA: piwi-interacting RNA
snoRNA: small nucleolar RNA
snRNA: small nuclear ribonucleic acid
tRNA: transfer RNA

## Declarations

### Ethics approval and consent to participate

Not applicable.

### Consent for publication

All authors have consented to the publication of this manuscript.

### Availability of data and material

These are available at https://regepi.bwh.harvard.edu/circurna/ and the source code is available at https://github.com/cougarlj/COMPASS.

### Authors’ contributions

JL built the platform COMPASS and wrote the manuscript. ATK evaluated COMPASS and exceRpt performances on the study data and revised the manuscript. RPC, LF and SSA prepared the performance study data. LP helped with the RNA annotation algorithm. KGT conceived the study and revised the manuscript. All authors have read and approved the final manuscript.

## Acknowledgements

The author wish to thank Jody Sylvia hosting COMPASS on the local test server.

## Additional files

Additional file 1. The comparisons of piRNAs, snRNAs, snoRNAs and tRNAs at the sample level.

