## Additional Files for "COMPASS: a COMprehensive Platform for smAll RNA-Seq data analySis"

**Additional file 1.** The comparisons of piRNAs, snRNAs, snoRNAs and tRNAs at the sample level.

**Table S1.** piRNAs identified by COMPASS and exceRpt among each sample.

|  | piRNA | | | | |
| --- | --- | --- | --- | --- | --- |
|  | COMPASS  (median) | exceRpt  (median) | Overlap | COMPASS_Unique | exceRpt_Unique |
| SAMPLE_1 | 35(686) | 18(569.83) | 16 | 19 | 2 |
| SAMPLE_2 | 71(892) | 26(2432.76) | 26 | 45 | 0 |
| SAMPLE_3 | 87(353) | 34(1020.79) | 33 | 54 | 1 |
| SAMPLE_4 | 64(503) | 25(1001.23) | 25 | 39 | 0 |
| SAMPLE_5 | 58(1134.5) | 28(771.13) | 27 | 31 | 1 |
| SAMPLE_6 | 56(1915.5) | 25(1771) | 22 | 34 | 3 |
| SAMPLE_7 | 84(675.5) | 44(507.13) | 42 | 42 | 2 |
| SAMPLE_8 | 55(925) | 26(1145.25) | 26 | 29 | 0 |
| SAMPLE_9 | 74(518.5) | 34(592.42) | 33 | 41 | 1 |
| SAMPLE_10 | 47(681) | 21(379) | 21 | 26 | 0 |
| SAMPLE_11 | 57(883) | 30(789.5) | 27 | 30 | 3 |
| SAMPLE_12 | 55(455) | 24(760.21) | 24 | 31 | 0 |

**Table S2.** snRNAs identified by COMPASS and exceRpt among each sample.

|  | snRNA | | | | |
| --- | --- | --- | --- | --- | --- |
|  | COMPASS  (median) | exceRpt  (median) | Overlap | COMPASS_Unique | exceRpt_Unique |
| SAMPLE_1 | 60(3974) | 46(296.30) | 43 | 17 | 3 |
| SAMPLE_2 | 112(358.5) | 82(82.84) | 71 | 41 | 11 |
| SAMPLE_3 | 123(130) | 64(342.40) | 59 | 64 | 5 |
| SAMPLE_4 | 87(638) | 57(295.49) | 52 | 35 | 5 |
| SAMPLE_5 | 43(744) | 36(22.41) | 32 | 11 | 4 |
| SAMPLE_6 | 47(1600) | 31(55.38) | 28 | 19 | 3 |
| SAMPLE_7 | 71(1098) | 54(118.71) | 48 | 23 | 6 |
| SAMPLE_8 | 95(198) | 87(22.21) | 80 | 15 | 7 |
| SAMPLE_9 | 100(117) | 52(148.87) | 48 | 52 | 4 |
| SAMPLE_10 | 57(834) | 34(45.11) | 28 | 29 | 6 |
| SAMPLE_11 | 59(1271) | 53(117.92) | 47 | 12 | 6 |
| SAMPLE_12 | 55(703) | 48(154.09) | 45 | 10 | 3 |

**Table S3.** snoRNAs identified by COMPASS and exceRpt among each sample.

|  | snoRNA | | | | |
| --- | --- | --- | --- | --- | --- |
|  | COMPASS  (median) | exceRpt  (median) | Overlap | COMPASS_Unique | exceRpt_Unique |
| SAMPLE_1 | 11(926) | 7(145.8) | 6 | 5 | 1 |
| SAMPLE_2 | 14(998) | 12(206) | 8 | 6 | 4 |
| SAMPLE_3 | 28(155) | 12(203.1) | 9 | 19 | 3 |
| SAMPLE_4 | 21(320) | 14(123.5) | 12 | 9 | 2 |
| SAMPLE_5 | 12(444) | 8(43.5) | 5 | 7 | 3 |
| SAMPLE_6 | 17(430) | 6(185.9) | 5 | 12 | 1 |
| SAMPLE_7 | 42(154) | 23(112) | 13 | 29 | 10 |
| SAMPLE_8 | 12(849) | 11(148.1) | 8 | 4 | 3 |
| SAMPLE_9 | 39(140) | 39(6.49) | 11 | 28 | 28 |
| SAMPLE_10 | 3(184) | 0(NA) | 0 | 3 | 0 |
| SAMPLE_11 | 9(550) | 4(298) | 1 | 8 | 3 |
| SAMPLE_12 | 9(748) | 6(53.9) | 5 | 4 | 1 |

**Table S4.** tRNAs identified by COMPASS and exceRpt among each sample.

|  | tRNA | | | | |
| --- | --- | --- | --- | --- | --- |
|  | COMPASS  (median) | exceRpt  (median) | Overlap | COMPASS_Unique | exceRpt_Unique |
| SAMPLE_1 | 13(11832) | 10(1318) | 10 | 3 | 0 |
| SAMPLE_2 | 19(12996) | 14(1168) | 14 | 5 | 0 |
| SAMPLE_3 | 22(7807.5) | 18(696.5) | 17 | 5 | 1 |
| SAMPLE_4 | 20(8699.5) | 13(1072) | 13 | 7 | 0 |
| SAMPLE_5 | 18(5860.5) | 12(700.05) | 12 | 6 | 0 |
| SAMPLE_6 | 20(12415) | 13(1700) | 13 | 7 | 0 |
| SAMPLE_7 | 21(10909) | 18(908.5) | 18 | 3 | 0 |
| SAMPLE_8 | 18(7321.5) | 13(539) | 13 | 5 | 0 |
| SAMPLE_9 | 23(5244) | 18(796) | 18 | 5 | 0 |
| SAMPLE_10 | 14(10299) | 11(742) | 10 | 4 | 1 |
| SAMPLE_11 | 19(7166) | 13(962) | 13 | 6 | 0 |
| SAMPLE_12 | 18(10052) | 12(684.5) | 11 | 7 | 1 |
